# Modulation of the spontaneous brain activity and functional connectivity in the triple resting-state networks following the visual oddball paradigm

**DOI:** 10.1101/2021.01.26.428223

**Authors:** Hasan Sbaihat, Ravichandran Rajkumar, Shukti Ramkiran, Abed Al-Nasser Assi, N. Jon Shah, Tanja Veselinović, Irene Neuner

## Abstract

The default mode network (DMN), the salience network (SN), and the central executive network (CEN) could be considered as the core resting-state brain networks (RSN) due to their involvement in a wide range of cognitive tasks. Despite the large body of knowledge relating to their regional spontaneous activity (RSA) and functional connectivity (FC) of these networks, less is known about the influence of task-associated activity on these parameters and on the interaction between these three networks. We have investigated the effects of the visual-oddball paradigm on three fMRI measures (amplitude of low-frequency fluctuations for RSA, regional homogeneity for local FC, and degree centrality for global FC) in these three core RSN networks. A rest-task-rest paradigm was used and the RSNs were identified using independent component analysis (ICA) on the resting-state data. We found that the task-related brain activity induced different patterns of significant changes within the three RS networks. Most changes were strongly associated with the task performance. Furthermore, the task-activity significantly increased the inter-network correlations between the SN and CEN as well as between the DMN and CEN, but not between the DMN and SN. A significant dynamical change in RSA, alongside local and global FC within the three core resting-state networks following a simple cognitive activity may be an expression of the distinct involvement of these networks in the performance of the task and their various outcomes.

## Introduction

Examination of regional spontaneous brain activity (RSA) and functional connectivity (FC) during resting-state (RS) conditions appears to be a promising approach for understanding brain organization at the systems level [1]. Within the several stable RS networks identified up to now, three networks stand out for their importance and synchronized interplay: the default mode network (DMN), the salience network (SN), and the central executive network (CEN). These networks are often jointly referred to as the triple network model [2] and are considered to be the core neurocognitive networks due to their involvement in a wide range of cognitive tasks [1,3,4].

Specifically, the DMN is known to be a task-negative network associated with self-referential thoughts and mind-wandering [5]. It shows decreased activation during tasks in which self-referential and stimulus-independent intellectual activity is not involved [6,7]. Even more, numerous studies have demonstrated that midline DMN regions are among the most efficiently wired brain areas, serving as global hubs that bridge different functional systems across the brain [8,9]. Increased DMN connectivity with regions of other brain networks has been shown to facilitate performance during goal-directed tasks [10]. Thus, DMN is not engaged only under resting-state conditions but also under task performance and post-task processes as well [10–12].

The CEN is a task-positive network, engaged in higher-order cognitive and attention control as well as in working memory, decision making and goal-directed behavior [13–15]. Conversely, the SN is involved in detecting, filtering and integrating relevant internal (e.g., autonomic input) and external (e.g., emotional information) salient stimuli in order to guide behavior [1,16]. Furthermore, it displays a crucial role in the functional and dynamic switching between the DMN and CEN (i.e., between task-based and task-free states) [17,18].

Dynamic interactions between the three networks of the triple network model influence cognition and emotion, affecting performance and impulsivity [19–21]. Moreover, an altered interaction between these networks has been shown in patients with major depressive disorder [22], post-traumatic stress disorder [23], obsessive-compulsive disorder [24], and schizophrenia [25,26]· Altogether, an increasing body of evidence suggests that aberrant function of the triple networks underlies the psychopathology of all major psychiatric disorders [27] and disturbed functional interactions among them may be considered a potential neurophysiological biomarker for different psychopathological phenomena across several neuropsychiatric disorders [28]. It is therefore particularly important to understand the physiological fluctuations in the activity and interactions of these networks in order to be able to differentiate them from pathological conditions.

Continuous fluctuations of the main properties of the networks (as RSA and FC) have been shown during rest and during task-associated activities [29,30]. Much less is known about the extent to which these properties can be influenced by a specific task and to what extent a task-associated activity affects the interaction between the networks.

A simple method to investigate the effects of task-related activation on the RSA is the rest-task-rest paradigm (RTR) [5,31]. To date, a task-induced modulation of the RSA has been observed following cognitive tasks involving working memory, emotion, visual perception, and motor training. However, previous studies have mainly focused on whole-brain [31–35] or on specific brain structures known to be involved in the tasks [36,37]. None of the mentioned studies has specifically addressed the impact of a task on the triple network. Moreover, previous investigations have overall changes in static connectivity in different time periods (before and after the task), but changes in the relationship between the different networks (particularly in the triple network, which is the focus of this study) remain poorly understood. Thus, in this study, we have specifically examined task-induced changes in RSA and FC in the triple network of the RS (DMN, SN and CEN) and the task-induced effects on the interactions between them.

Concretely, this study aims to assess the extent of the influence exerted by a well-established task - the visual oddball paradigm [38] on the post-task RS in the regions of the triple network using the RTR design. The visual oddball paradigm task was chosen as it elicits the blood oxygen level dependent (BOLD) response in a large set of distributed networks [39–43]. In particular, the task performance is associated with activation in brain regions linked to the three networks (the SN [44], the dorsolateral prefrontal cortex (CEN) [45,46], and the cingulate and prefrontal cortex (DMN) [47].

For the identification of the triple network regions we applied a group independent component analysis (ICA) to the RS data. Several different measures of FC can be calculated from fMRI, each reflecting a different property of the brain networks. For this approach, we chose two such measures, the regional homogeneity (ReHo) [48] the degree centrality (DC) [49], as these are suitable for investigating the voxel level local and global FC, respectively. Furthermore, the amplitude of low-frequency fluctuations (ALFF) [50], is suitable for depicting the RSA. Combining these measures enables the complementary characterization of changes in activation and communication of specific networks or regions.

We hypothesised that the task-based activity would distinctly affect the RS RSA as well as the local and the global connectivity in the triple network. Due to the central role of the SN during the occurrence of salient stimuli or during the performance of a cognitive task, we also expected internetwork functional connectivity to increase between the SN and the other two networks of the triple network model (DMN, CEN).

## Materials and Methods

### Subjects

21 right-handed healthy subjects (17 males and four females) were included in this study (age range between 19 to 40 years; mean: 29 ± 5.6 years). All subjects were healthy and without a history of neurological or psychiatric disorders. The study was approved by the Ethics Committee of the Medical Faculty of the RWTH Aachen University, Germany. Written informed consent was obtained from all subjects following the recommendations of the Declaration of Helsinki.

### Experimental design

To investigate the effects of task-induced brain activity on the post-task resting-state, the experiment followed a rest-task-rest (RTR) design consisting of three parts – each part representing a different brain state: first RS (R1), active state (during the performance of the visual oddball paradigm) and the second, post-task RS (R2) (Fig 1).

**Fig 1.**
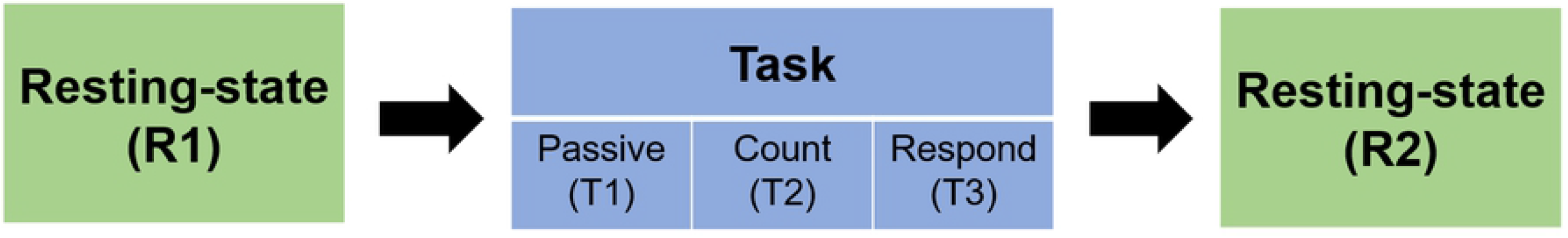
Experimental design of the rest-task-rest paradigm (RTR) which includes two resting-state conditions (pre- and post-task resting-state, R1 and R2) and the task condition composed of three subtasks of the visual oddball paradigm (VOP).

During RS conditions, the subjects were instructed to close their eyes and not focus on any specific thoughts. All the fMRI data were acquired in a single scanning session and instructions were given to the subjects in-between each condition via a microphone.

The visual oddball paradigm comprises of three subtasks: passive (T1), count (T2), and respond (T3). Two different colored circles were established as frequent (yellow circles) and target (blue circles) stimuli. During the passive condition, the subjects were asked to simply keep the stimuli under observation. During the count condition, the subjects were asked to count the target stimuli (blue circles), and during the respond condition, the subjects were instructed to press a button with their right index finger as soon as they recognised the target stimuli.

Each visual oddball paradigm condition included 200 trials (160 frequent and 40 target stimuli). The single stimulus was 30 cm in diameter shown on a black background for 500 milliseconds with a variable interstimulus interval (ISI) of 500–10,000 milliseconds. The stimulus generator board (ViSaGe MKII, Cambridge Research System Ltd.) was used to generate the stimuli and a thin-film transistor display was used to view the stimuli. The thin-film transistor display was installed behind the scanner and was viewed using a mirror placed on the head coil of the magnetic resonance (MR) scanner.

A part of this data set (N = 16), which mainly focused on the analysis of the effects of different response modalities on the fMRI BOLD activation during the visual oddball paradigm, has been published previously [51].

### MR Data Acquisition

MR data were acquired using a 3T scanner (TIM-Trio, Siemens Healthineers, Erlangen, Germany). Sponge pads were used to reduce motion artefacts by limiting the subject’s head movement. The fMRI data were acquired using an echo planner imaging (EPI) sequence. The number of volumes were 304 for each task and 180 for each RS condition (repetition time (TR) = 2000 ms, echo time (TE) = 30 ms, flip angle (FA) = 79°, field of view (FOV) = 200 × 200 mm, 64 × 64 matrix, slice thickness = 3 mm, number of slices = 33).

Structural images were acquired using a magnetization prepared rapid gradient echo (MP-RAGE) sequence (TR = 2250 ms, TE = 3.03 ms, FA = 9°, FOV = 256 × 265 mm, 64 × 64 matrix, 176 slices, voxel size 1 × 1 × 1 mm^3^).

### fMRI Data Analysis

#### Task Data

The analysis of the task-related brain activation was performed using FSL software package (FMRIB’s Software Library, www.fmrib.ox.ac.uk/fsl). The pre-processing included slice timing correction, brain extraction (using BET) [52], motion correction (MCFLIRT) [53], spatial smoothing using a Gaussian kernel of full width at half maximum (FWHM) of 5 mm, and high pass temporal filtering (100s). A time-series of BOLD signal based on the general linear modal for each individual data set was performed using FILM with local autocorrelation correction [54]. The functional images were registered to the high-resolution structural images and subsequently to the Montreal Neurological Institute (MNI) standard space using the FLIRT tool [55]. The first-level analysis was performed with two explanatory variables (EV). The EVs were convolved with a double-gamma hemodynamic response function (HRF). Four contrasts were then created: target stimuli, frequent stimuli, target > frequent, frequent > target.

Group-level mixed-effects analysis was performed for the passive, count and respond sub-tasks to create a mean for each first level contrast using FLAME with spatial normalization to MNI space and using a cluster with a significance threshold of Z > 2.3, p = 0.05 [56]. A tripled two-group difference (“tripled t-test”) Was performed to evaluate the additional activation added to the passive condition by the count and respond conditions. The activation pattern regions were defined using Harvard-Oxford Cortical Structural Atlas in FSL software (FMRIB, Oxford, UK).

### Triple network identification

The multivariate exploratory linear decomposition into independent components (MELODIC) tool from the FSL software package was used to identify the triple networks (DMN, CEN, and SN) using pre-task RS fMRI data. Subject level RS-fMRI data were pre-processed as follows: the first eight fMRI volume images were removed, followed by slice timing correction, brain extraction (BET) [52], motion correction (MCFLIRT) [53], spatial smoothing FWHM = 5 mm, and high-pass temporal filtering 125s. The functional MRI images were co-registered linearly to high-resolution structural images and nonlinearly to MNI standard space using FLIRT [55]. Group ICA analysis was used to decompose the pre-task RS data into 20 components.

To identify the triple networks, a cross-correlation was performed between the functional brain networks atlas [57] and each of the ICA components. The cross-correlation was performed using the FSLUTILS (https://fsl.fmrib.ox.ac.uk/fsl/fslwiki/Fslutils) tool implemented in the FSL software package. ICA components that showed maximum correlation with each of the three networks in the functional brain networks atlas were chosen. The identified brain networks were binarized and used in the subsequent analysis as masks. The binarized masks were corrected for grey matter (GM) by including the voxels which showed more than 50% probability of being GM. The GM correction was performed using a tissue segmented MNI152 (2 × 2 × 2 mm^3^) template.

### fMRI measures calculation

The fMRI measures were computed for both the tasks and RS-fMRI using data processing and analysis for brain imaging (DPABI) [58], and SPM12 (http://www.fil.ion.ucl.ac.uk/spm/) toolboxes built on MATLAB software package version 2017b (The Math Works, Inc., Natick, MA, USA). Pre-processing was performed using the data processing assistant for the RS-fMRI (DPARSF) [59] advanced edition as follows: first eight fMRI volume images of each condition in each subject’s dataset were removed, followed by slice timing correction, realignment, nuisance covariates regression (NCR) and temporal filtering between 0.01 and 0.08 Hz. To get rid of the nuisance signals, the Friston 24-parameter model was used for covariate regression. The fMRI measures were calculated for each subject separately in individual brain imaging space. The DC was computed by applying a Pearson correlation coefficient between the time series of a given voxel and all other voxels in the whole brain by thresholding each correlation at (r > 0.25, p ≤ 0.001) [60]. ReHo was calculated by estimating the synchronization or similarity between the time series of a given voxel and 26 nearby neighbor voxels [48] using Kendall’s coefficient of concordance (KCC) [61]. The ALFF was calculated within the low-frequency range (0.01 – 0.1 Hz) [62]. The fMRI measures were normalised using a Z-value standardization procedure by subtracting the mean from each voxel and then dividing the value by the standard deviation of the whole brain. The Z-value standardised measures were co-registered to the MNI standard space (2 × 2× 2mm^3^), and, finally, spatial smoothing with FWHM at 4mm^3^ was performed.

### Further calculated values

The fMRI measures ALFF, ReHo, and DC were extracted from all voxels of the triple network for each condition in all subjects using the binarized triple network masks. The extracted voxel-level values were used to calculate several parameters of interest, relevant for the examination of the task effect on the post task resting-state. These parameters and the exact description of how they were calculated are shown in Table 1.

**Table 1:** Description of parameters used to examine the effect of the task on the fMRI measures in the post-task RS.

| Parameter | Calculation procedure/ meaning |
| --- | --- |
| **R1** | Voxel-level fMRI measures during the first (pre-task) resting-state (RS) (baseline) |
| **R2** | Voxel-level fMRI measures during the second (post-task) RS. |
| **T1, T2, T3** | Voxel-level fMRI measures during the three subtasks of the visual oddball paradigm. |
| **RS difference (RSD)** | Difference between post- and pre-task RS ($R2 - R1$) in the voxel-level fMRI measures for each subject. |
| **Task<sub>(whole)</sub>** | $\text{Task}_{(whole)} = (T1 + T2 + T3) / 3$<br>(mean values of the fMRI measures during the three subtasks) |
| **Main task<sub>(whole)</sub>** | $\text{Task}_{(whole)} - R1$ |
| **RS similarity (RSS)** | Correlation coefficient between R1 and R2 for each subject. |
| **Task effect at the group level** | Correlation coefficients between the differences ($\text{Task}_{(whole)} - R1$) and ($R2 - R1$).<br>All correlation coefficients were computed using Pearson's correlation coefficients at a significance level of $p < 0.05$. |

The inter-network FC of the three networks were calculated by first extracting the mean of the BOLD signal time series from the binarized mask of each network, followed by the computation of the Pearson’s correlation coefficient between each pair of networks. Fisher r to z transformation was performed to improve the normal distribution. A paired t-test was used to examine the difference of FC between the pre- and the post-task RS.

To investigate the relationship between the behavioral data (e.g. reaction time) and the fMRI measures, the correlation coefficients between the RSD and subject’s reaction time in the response condition was performed. Having checked the normality of the data using the Kolmogorov-Smirnov test, a paired-sample t-test was used in order to find the differences between the pre- and post-task RS in each fMRI measure.

## Results

### Behavioural data

The mean reaction time of the respond condition was 477ms (SD = 13).

### Imaging data - task data

The task data were initially analysed and reported following the examination of the first 16 participants [51]. The current analysis includes an enlarged collective of test subjects (N = 21) and confirms previously reported findings. In summary, activation in regions associated with a response to visual stimulation (occipital cortex) for both the target and the frequent stimuli was observed during all three subtasks of the visual oddball paradigm. Both, the count and respond conditions differed significantly from the passive condition in a number of brain regions including the pre- and post-central gyri, regions of the parietal cortex and the middle and inferior frontal gyri. Compared to the count condition, the response contrast yielded significant differences in the parietal operculum, inferior parietal lobule, insula, anterior cingulate cortex, and the posterior cingulate cortex (PCC).

### Imaging data -Triple network resting-state data

The triple network was identified using group independent component analysis (Fig 2). Specifically, the DMN included the posterior cingulate cortex (PCC), precuneus, angular gyrus, and medial prefrontal cortex (mPFC); the CEN included the lateral posterior parietal cortex (LPPC) and dorsolateral prefrontal cortex (DLPFC); the SN included the frontal insular cortex (FIC), and anterior cingulate cortex (ACC).

**Fig 2.**
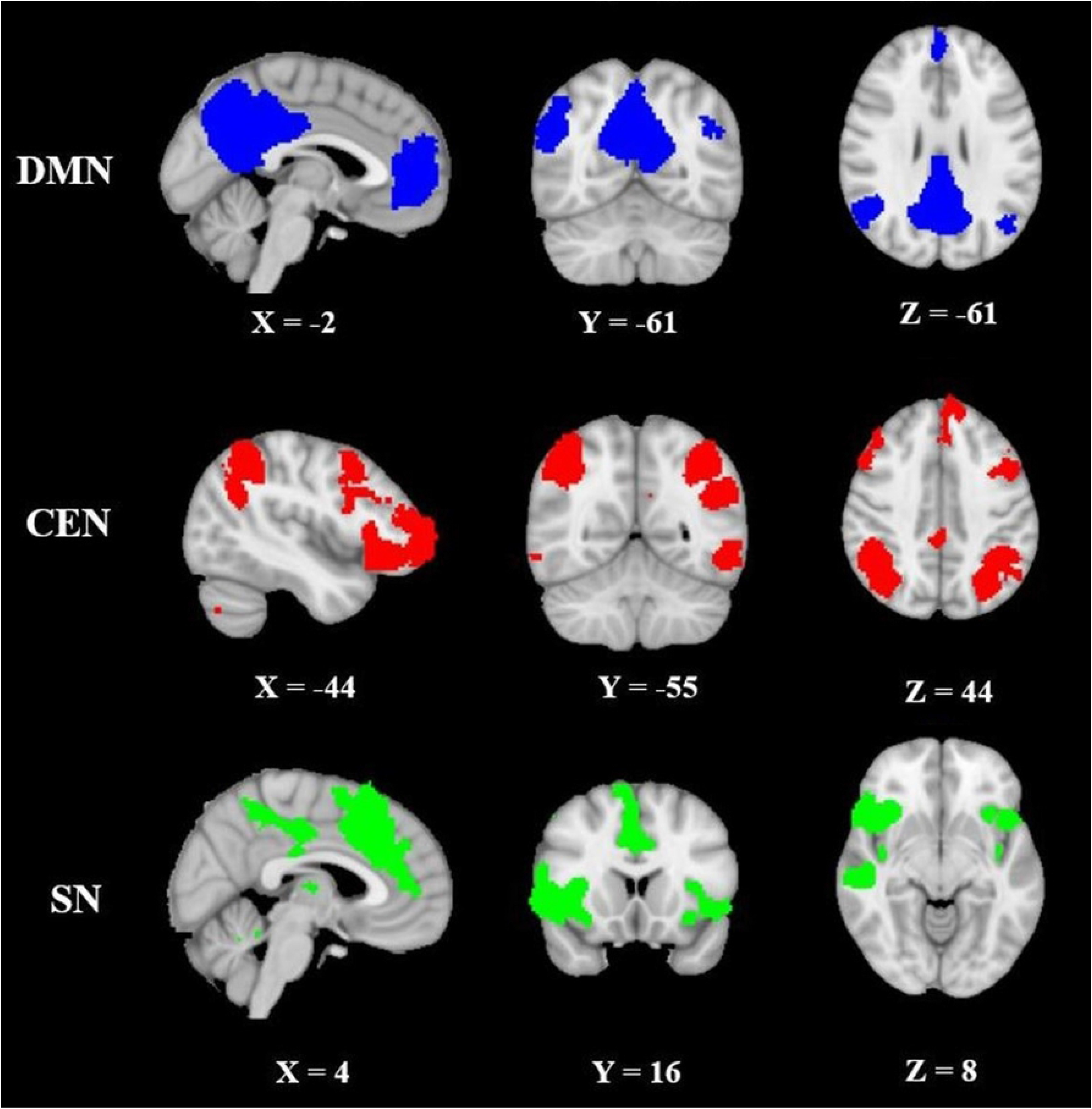
Depiction of the triple networks referred to as the triple network: default mode network (DMN, blue colour), central executive network (CEN, red colour), and salience network (SN, green colour). The networks were identified by decomposing the pre-task resting-state condition into 20 components from 21 subjects.

### RSA and FC across different brain-states

The fMRI measures showed different values in the RSA and the local and global FC during the different brain-states (rest-task-rest) (Fig 3). These values differed significantly at the group level. A pairwise comparison between the pre- and post-task RS (R1 and R2) revealed significant differences in each network and for each fMRI measure at a significance level of p < 0.05, with exception of the ReHo measure in the SN (Table 2).

**Fig 3.**
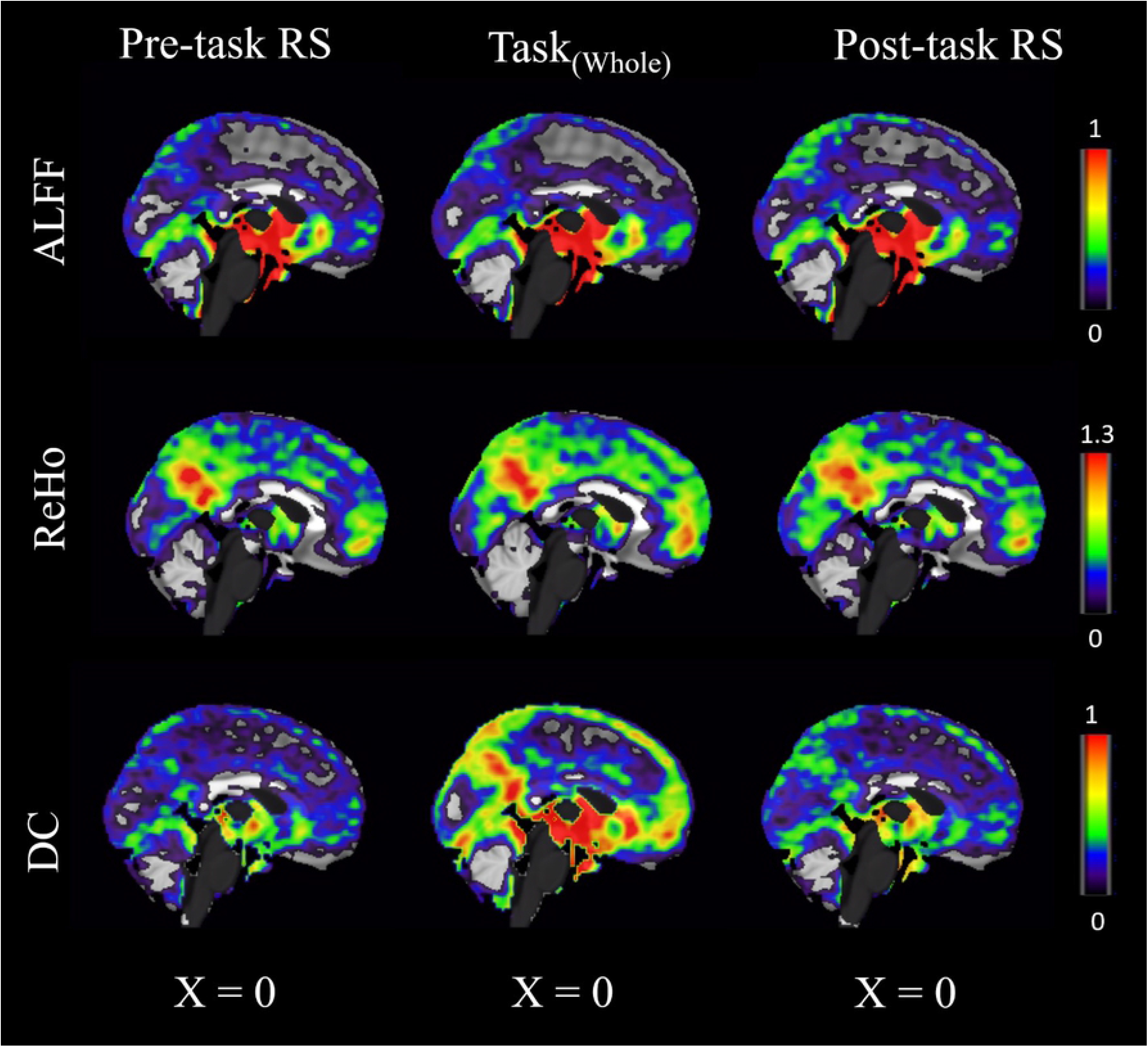
fMRI measures (ALFF, ReHo, and DC) from 21 subjects. Visual inspection shows a change in each of the three fMRI measures in the post-task resting-state (R2) when compared to pre-task resting-state networks (R1). The differences between R1 and R2 were significant in each network and in each fMRI measure at significance level p < 0.05, with exception of the ReHo measure in the salience network (SN).

**Table 2.**
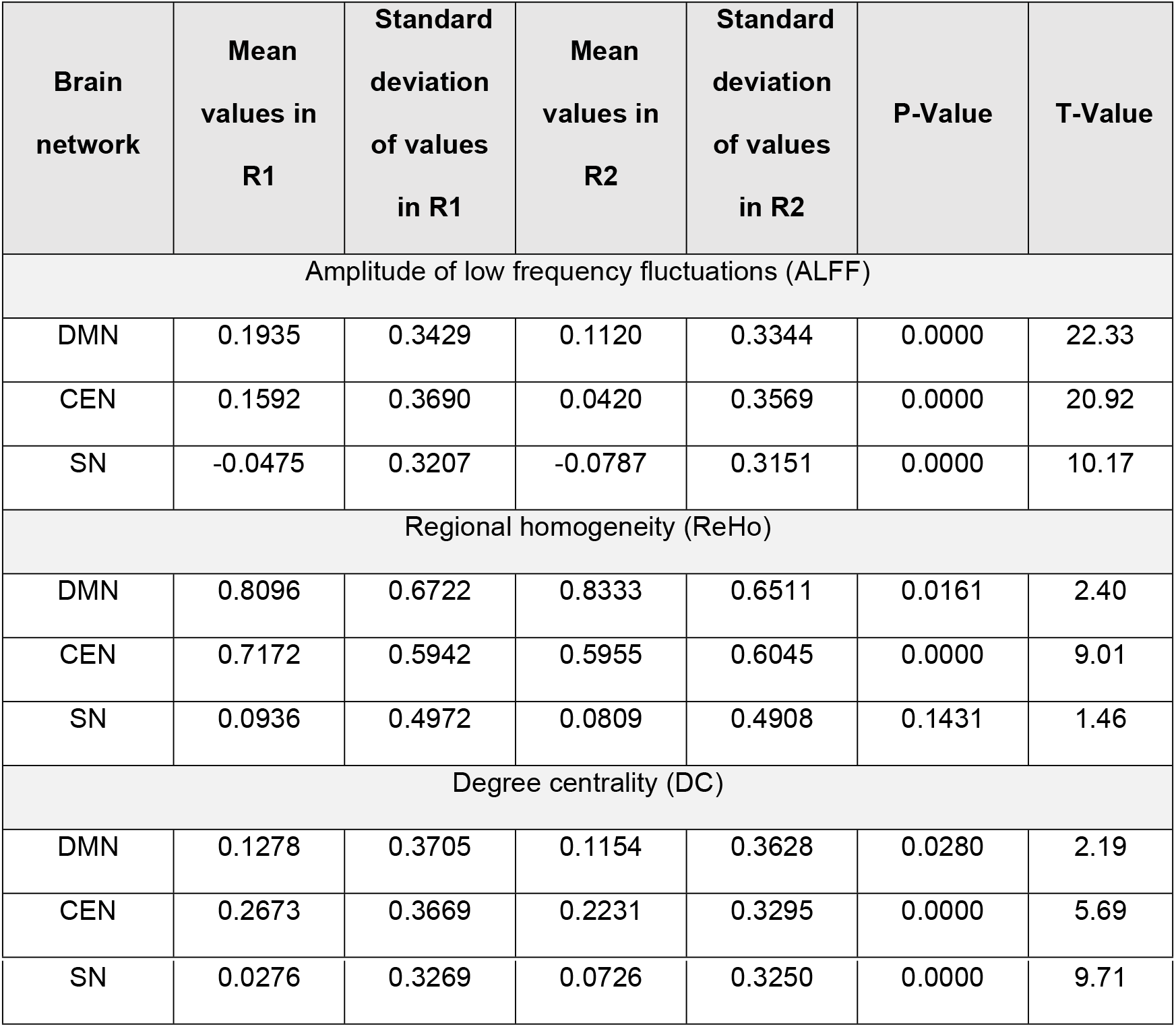
A paired-samples t-test was performed to compare the pre-task resting-state (R1) and post-task resting-state (R2) conditions in the triple network (default mode network (DMN), salience network (SN), and central executive network (CEN)), in each fMRI measurement (21 Subjects). There was a significant difference of p < 0.05 between R1 and R2 in each network and in each fMRI measurement, with the exception of the SN from the ReHo measurement.

| Brain network | Mean values in R1 | Standard deviation of values in R1 | Mean values in R2 | Standard deviation of values in R2 | P-Value | T-Value |
| --- | --- | --- | --- | --- | --- | --- |
| Amplitude of low frequency fluctuations (ALFF) |  |  |  |  |  |  |
| DMN | 0.1935 | 0.3429 | 0.1120 | 0.3344 | 0.0000 | 22.33 |
| CEN | 0.1592 | 0.3690 | 0.0420 | 0.3569 | 0.0000 | 20.92 |
| SN | -0.0475 | 0.3207 | -0.0787 | 0.3151 | 0.0000 | 10.17 |
| Regional homogeneity (ReHo) |  |  |  |  |  |  |
| DMN | 0.8096 | 0.6722 | 0.8333 | 0.6511 | 0.0161 | 2.40 |
| CEN | 0.7172 | 0.5942 | 0.5955 | 0.6045 | 0.0000 | 9.01 |
| SN | 0.0936 | 0.4972 | 0.0809 | 0.4908 | 0.1431 | 1.46 |
| Degree centrality (DC) |  |  |  |  |  |  |
| DMN | 0.1278 | 0.3705 | 0.1154 | 0.3628 | 0.0280 | 2.19 |
| CEN | 0.2673 | 0.3669 | 0.2231 | 0.3295 | 0.0000 | 5.69 |
| SN | 0.0276 | 0.3269 | 0.0726 | 0.3250 | 0.0000 | 9.71 |

Concretely, the ALFF values decreased significantly in all three observed networks (p < 0.001 in all three networks). For the local connectivity parameter (ReHo), a significant increase in the DMN (p = 0.016) was observed, while the ReHo value decreased in the CEN (p < 0.001) and remained without statistically significant alteration in the SN. The long-range connectivity (DC) decreased significantly in the DMN and CEN (p = 0.028; p < 0.001), while increasing in the SN (p < 0.001).

### Associations between the pre- and post-task resting-state differences and the task

The RSS values were calculated separately for the triple networks (DMN, CEN and SN) for each of the fMRI measures (ALFF; ReHo and DC) are shown in Table 3. The RSS values were comparable for all three parameters across all three networks.

**Table 3.**
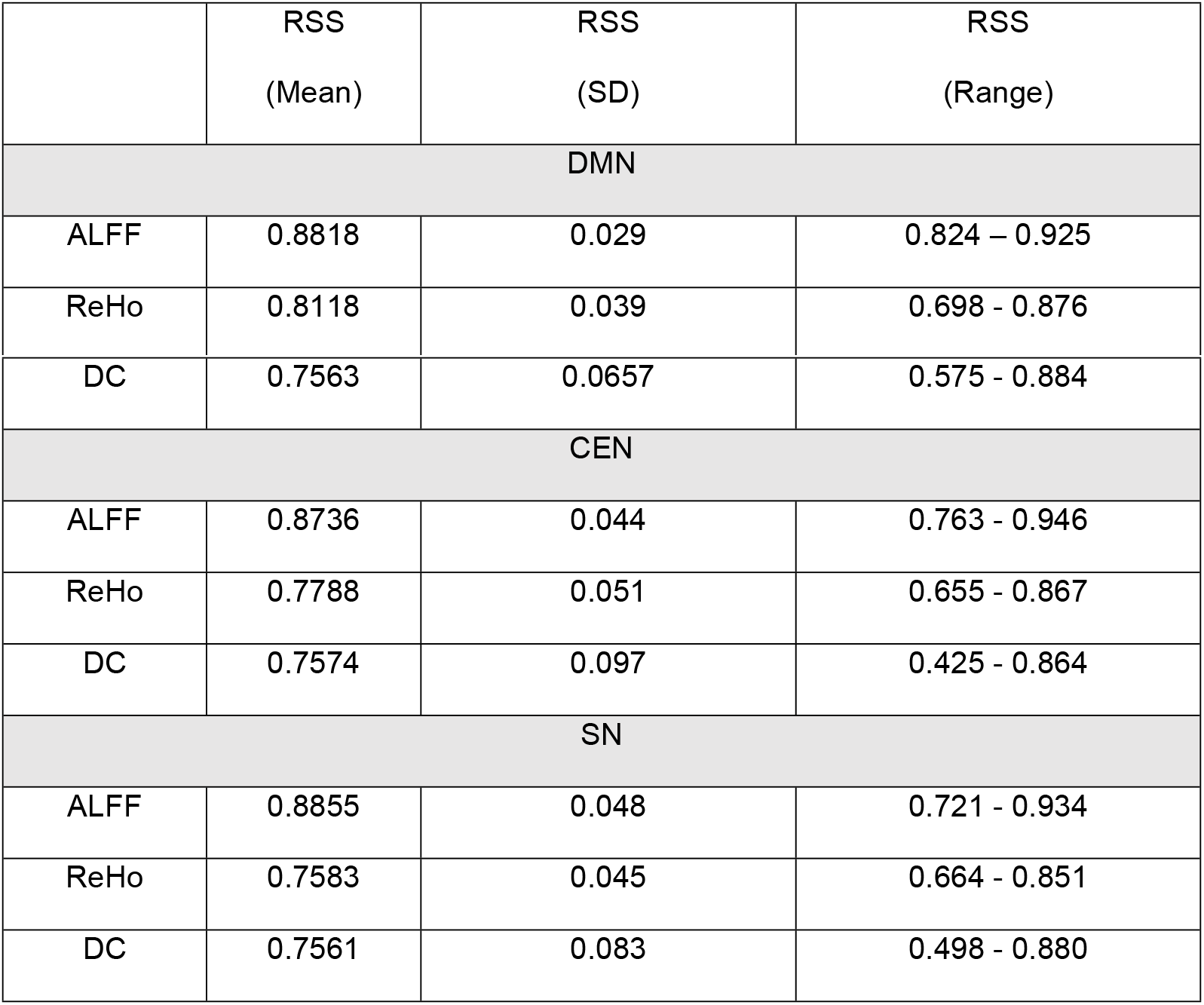
Mean values, standard deviation, and the range of the resting-state similarity (RSS) calculated separately for each resting-state fMRI parameter (ALFF, ReHo, and DC) and for each of the triple networks (default mode network (DMN), salience network (SN), and central executive network (CEN).

|  | RSS<br>(Mean) | RSS<br>(SD) | RSS<br>(Range) |
| --- | --- | --- | --- |
| DMN |  |  |  |
| ALFF | 0.8818 | 0.029 | 0.824 – 0.925 |
| ReHo | 0.8118 | 0.039 | 0.698 - 0.876 |
| DC | 0.7563 | 0.0657 | 0.575 - 0.884 |
| CEN |  |  |  |
| ALFF | 0.8736 | 0.044 | 0.763 - 0.946 |
| ReHo | 0.7788 | 0.051 | 0.655 - 0.867 |
| DC | 0.7574 | 0.097 | 0.425 - 0.864 |
| SN |  |  |  |
| ALFF | 0.8855 | 0.048 | 0.721 - 0.934 |
| ReHo | 0.7583 | 0.045 | 0.664 - 0.851 |
| DC | 0.7561 | 0.083 | 0.498 - 0.880 |

The correlation between the differences between post-task and pre-task RS parameters (RSD = R2 - R1) and the fMRI measures resulting from the pure task effects (task_(Whole)_ - R1) are depicted in (Fig 4). Significant positive correlations were found in DMN for ALFF (r = 0.48, p = 0.02) and DC (r = 0.58, p = 0.005); in CEN for ALFF (r = 0.44, p = 0.04), ReHo (r = 0.69, p = 0.004) and DC (r = 0.67, p = 0.008); and in SN for ALFF (r = 0.69, p = 0.004), ReHo (r = 0.58, p = 0.004), and DC (r = 0.49, p = 0.02).

**Fig 4.**
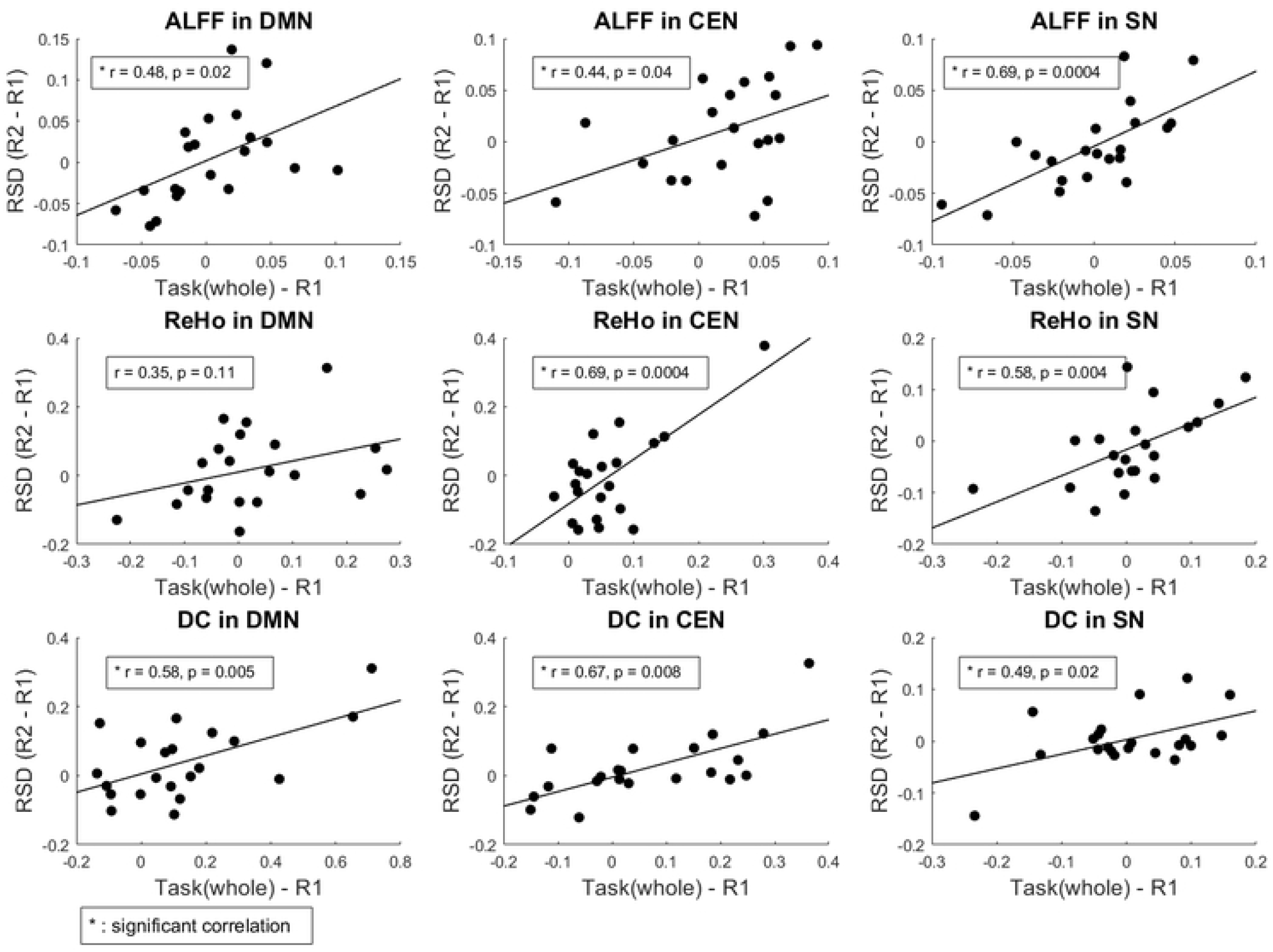
Correlations between the fMRI measures resulting from the pure task_(whole)_ effects and the RSD in the triple network, including the DMN, CEN, and SN of the fMRI measures.

Further, a significant negative correlation was observed between the RS differences (RSD) in DC and the subject’s reaction time in the respond condition in the SN (r = -0.46, p = 0.04), but not in ReHo or ALFF measurements. No significant correlations between the fMRI measurements and the reaction time could be observed in DMN and CEN (Fig 5).

**Fig 5.**
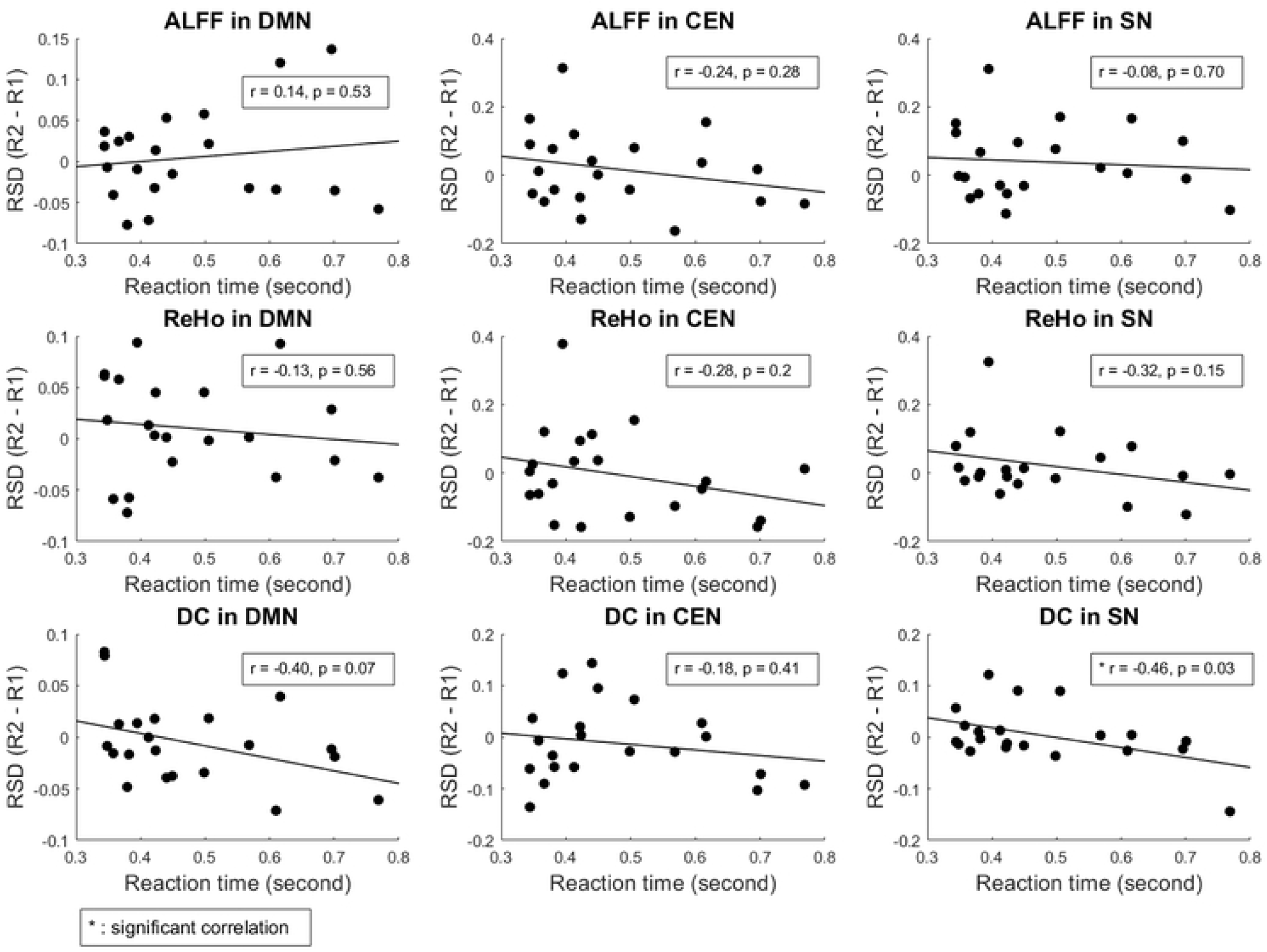
Correlation between the resting-state differences (RSD) and the subject’s reaction time to the respond condition of the VOP, depicted in the triple networks and each fMRI measurement. Only the SN shows a significant negative correlation in the DC measurement.

### Inter-network interaction

The functional connectivity between the DMN and CEN increased significantly following the performance of the task (p = 0.015). The connectivity strength between the DMN and the SN remained stable (p = 0.25), whereas it increased significantly between the SN and CEN (p = 0.0004) (Fig 6).

**Fig 6.**
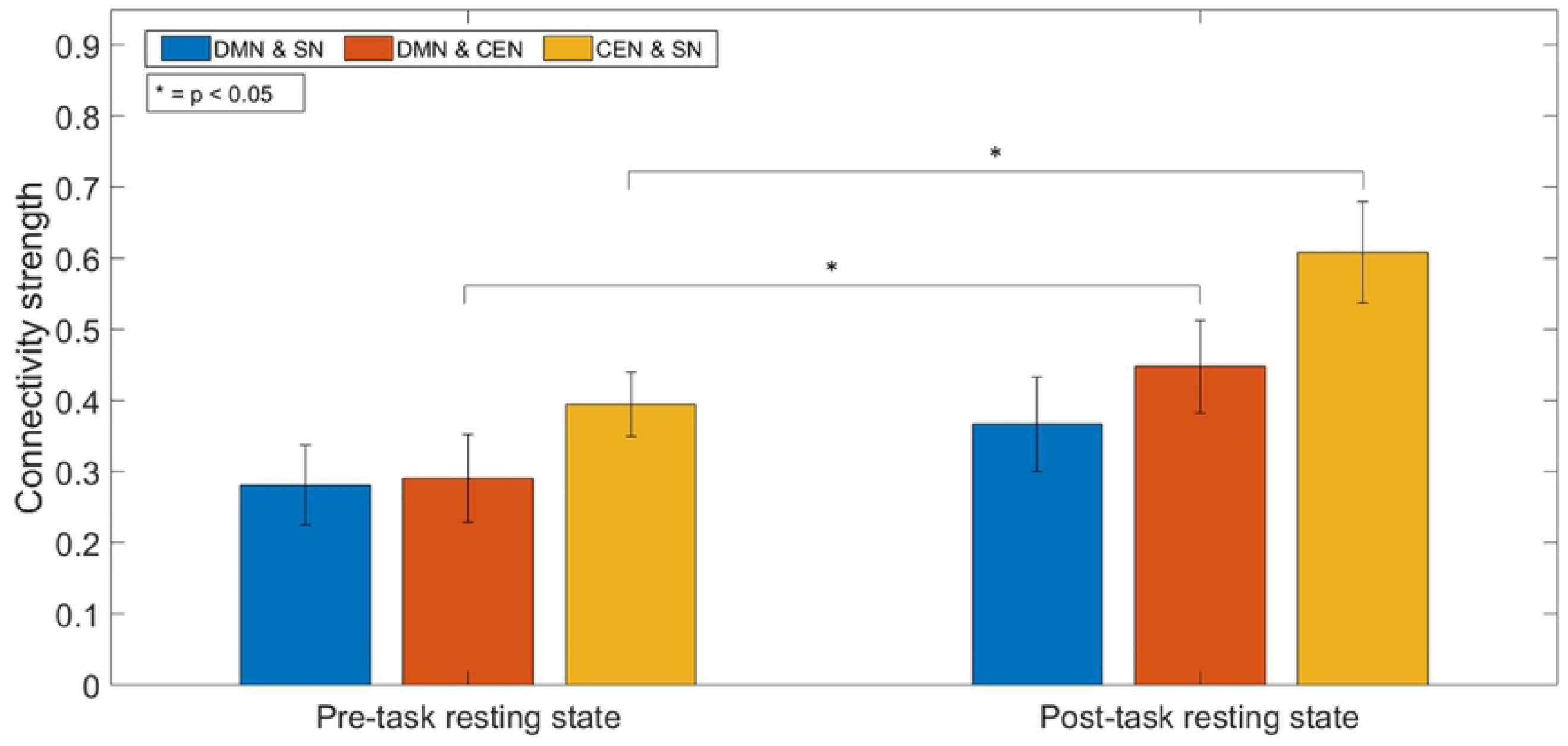
Strength of the FC between each pair of networks in the triple network in the pre- and post-task resting-state. There is a significant increase in FC between the DMN and CEN, and between the CEN and the SN in the post-task resting-state (p < 0.05). The bars represent the standard error.

## Discussion

In this study, we investigated the effects of a simple visual-oddball paradigm on three basic fMRI measurements of the RS – ALFF (RSA), ReHo, and DC (the local and the global functional connectivity, respectively) - in the three networks - DMN, CEN, and SN. Our analysis revealed that the brain activity following completion of the task had a significant effect on all examined parameters in all networks, except for the measure of local connectivity (ReHo) in the SN. Furthermore, the task performance induced a significant increase in the inter-network correlations between the SN and CEN, as well as between the DMN and CEN, but not between the DMN and SN. Also, the differences between the pre- and the post-task RS (R2 - R1) were strongly associated with the main task influence (task_(Whole)_ - R1) in all three networks (ALFF and DC in the three networks, ReHo in the CEN and SN). Finally, at a behavioral level, the task performance (subject’s reaction time in the respond condition) correlated solely with the RS difference in DC for the SN.

Our findings indicate dynamic, disparate alterations in the post-task resting-state brain networks as a function of immediately preceding cognitive experiences. Thereby, the extent of the changes in the RS networks can be said to be closely associated with the magnitude of the direct task-effects measurable during the task performance.

Within the DMN, the ALFF values decreased significantly, indicating a reduction of the DMN RSA following task performance. This effect has been reported previously [63–65]. As the DMN is regularly deactivated during the task performance [66,67], the continued reduction of the RSA observed after the task could be an expression of a redistribution of cognitive resources in the subsequent rest phase but could also indicate a neuronal correlate of task-induced temporal fatigue after a cognitive engagement [64]. In our study, the ALFF decrease in the DMN was accompanied by a decrease in DC. Similar findings have been previously reported after subjects performed a sustained auditory working memory task [65]. Interestingly, the local FC (ReHo) increased significantly in the DMN. Local FC is defined by the temporal coherence or synchronization of the BOLD time series within a set of a given voxel’s nearest neighbors [26]. ReHo represents the most efficient, reliable, and widely used index of local FC [68,69]. An increase in ReHo indicates an increased local synchronization of spontaneous neural activity [65]. Moreover, it was previously postulated that ReHo correlates with measures of functional segregation such as local efficiency and clustering coefficients [70]. Thus, increased ReHo in the DMN following task completion may reflect a restriction of information transfer to spatially close areas, as well as functional segregation from distant hubs and decreased communication with remote brain regions [71]. This result complements the observation of the decreased DC values in the post-task RS in DMN.

A significant increase in global brain connectivity (DC) was found in the SN in the post-task RS. This finding is in concordance with the established role of the SN as a network known to demonstrate competitive interactions during cognitive information processing [6,19] and, thus, having a critical role in switching between two other major RS networks (the DMN and the CEN [1]. In particular, the main hubs of the SN, the frontal inferior insula and ACC, are known to share significant topographic reciprocal connectivity and form a tightly coupled network, ideally placed to integrate information from several brain regions [72,73]. Thus, they seem to moderate arousal during cognitively demanding tasks and play a unique function in initiating control signals that activate the CEN and deactivate the DMN [74].

The finding of a significant decrease of the RSA (ALFF) in the post-task RS in SN is slightly more complex to explain. Previous investigations have linked increased ALFF values in some parts of the SN to a hyperarousal state in patients with MDD [75]. The reduction of the RSA in the post-task RS in our study may be an expression of a decreased arousal and decreased stimulus monitoring immediately after a completion of a task.

The connectivity analysis between the three networks revealed an increased synchronization (in terms of a significantly increased connectivity strength) for the SN with the CEN but not with the DMN in the post-task RS compared to the pre-task RS. This may be an after-task of the inter-network interactions during the paradigm performance. Indeed, Sridharan and colleagues have shown that the connectivity strength during the visual oddball paradigm particularly increased between the main nodes of the SN (frontal anterior insula and ACC) and all main nodes of the CEN, while the interactions between the SN and DMN were less pronounced [74].

In the CEN, the spontaneous brain activity (ALFF) decreased alongside the measures of the local and global connectivity. At a broad level, the CEN is included in higher order executive functioning, including the cognitive control of thought, emotion regulation, and working memory [16,76,77] and is thus activated during efforts to exert self-control, reappraise threatening stimuli, and to suppress intrusive, unpleasant thoughts [78–80]. CEN activity has been shown to be anti-correlated with activity in the DMN in healthy adults [1,19,74], while some investigations indicate that the CEN also exhibits an inhibitory control on the DMN [81]. Thus, the decrease in RSA in the CEN following completion of a cognitive paradigm may be the basis for the restoration of the regular activity of the DMN within the scope of a decline in DMN inhibition which occurred as a result of increased CEN activity during the task performance.

Interestingly, the connectivity between the CEN and DMN also increased in the post-task resting-state. This finding is consistent with the literature on the cooperative activity of the DMN and the CEN during different mental operations [82]. An increased coupling between some parts of these two networks has been shown in problem-solving tasks [83], social working memory [84], and during creative idea production [85]. Furthermore, a significant interaction between the DMN and the CEN has also been shown during the RS condition [86]. Thereby, this interaction seems to fluctuate dynamically across short time scales [87], indicating that the temporal relationships between the DMN and CEN shifts depending on the change in the attention focus and the immediately preceding activity. Thus, the increased connectivity between the DMN and the CEN in the post-task RS observed in our study may be an expression of the shifting of attention after task completion.

Several subregions of the triple networks are known to be activated during the performance of cognitively demanding tasks [88]. In the case of the visual oddball paradigm performed in our study, the main task specific activation has been reported previously by Warbrick and colleagues [51]. The target detection specifically involved parts of the DMN (PCC) and the SN (Insula, ACC). The insula activation was common to the count and respond conditions. The intensive involvement of different subregions of the triple networks in the performance of the task may have contributed to the significant changes in the triple network model networks in the post-task RS compared to the pre-task RS. Indeed, we have found positive correlations between the extent of the differences between R1 and R2 regarding specific parameters and the actual task effect on the same parameters in the triple networks. These correlations were significant in the DMN for ALFF and DC measures and in the CEN and the SN for all three fMRI measures. A close relationship between the cognitive level of the previous task and the extent of the modulation in the brain networks has been reported previously. Barnes and colleagues observed that the changes in endogenous dynamics in post-task RS is directly related to the difficulty of task performance [89]. In the case of the visual oddball paradigm used here, the levels of cognitive demand for all the three subtasks are not widely different and the whole paradigm did not require high cognitive effort. However, we observed that the extent of the changes in the RSA and local as well as global connectivity in the core RS networks in the post-task condition follows the extent of the task-induced changes within those networks. Thus, the task-induced modification of the RS activity and connectivity seems to be influenced by the intensity of the immediately preceding activation within the observed regions/networks.

A significant correlation between the behavioral outcomes in the visual oddball paradigm and the changes in the fMRI parameters could only be observed in the SN. Participants showing better performance (shorter response times in the response subtask) had a higher increase in global connectivity when comparing the second and the first RS. Thus, higher flexibility of the SN may be associated with better cognitive performance. This supports the observation that subjects with lower SN-network interactions have more pronounced inattention scores [90].

## Conclusion

Our findings confirm significant dynamical changes in RSA, alongside local and global connectivity within the triple networks following a simple cognitive activity. As discussed above, the change in patterns differed noticeably between the networks and was tightly associated with the task-related brain activity. The observed changes may be an expression of the distinct involvement of the networks in the performance of the task and their various roles in the processing and integration of the immediately preceding experience. Our results provide further insight into the dynamics within and between the triple networks and contribute to a better understanding of their functional importance and interplay.

## Acknowledgements

This study is considered to be part of the doctoral thesis (Dr. rer. medic.) of Mr. Hasan Sbaihat, Faculty of Medicine, RWTH Aachen University, Germany. Hasan Sbaihat would like to thank the Palestinian German Science Bridge (PGSB), Federal Ministry of Education and Research Germany, and the Palestine Academy for Science and Technology (PALAST) for their assistance and scholarship funding. The authors would like to thank Dr. Jorge Arrubla and Tracy Warbrick for their help in data acquisition, and Dr. Shivakumar Viswanathan for providing advice on statistics. Finally, authors would like to thank Ms. Claire Rick and Mr. Dennis Thomas for proofreading the manuscript.

